# Differential Tractography as a Track-Based Biomarker for Neuronal Injury

**DOI:** 10.1101/576025

**Authors:** Fang-Cheng Yeh, Islam M. Zaydan, Valerie R. Suski, David Lacomis, R. Mark Richardson, Joseph C. Maroon, Jessica Barrios-Martinez

**Author notes:** Correspondence to: Fang-Cheng Yeh, M.D. Ph.D., Department of Neurological Surgery, Department of Bioengineering, University of Pittsburgh, Pittsburgh, Pennsylvania. **One Sentence Summary:** Differential tractography utilizes repeat diffusion MRI scans to identify the exact segment of tracks with a neuronal injury.

## Abstract

Diffusion MRI tractography has been used to map the axonal structure of human brain, but its ability to detect neuronal injury is yet to be explored. Here we report differential tractography, a new type of tractography that utilizes repeat MRI scans and a novel tracking strategy to map the exact segment of fiber pathways with a neuronal injury. We examined differential tractography on multiple sclerosis, Huntington disease, amyotrophic lateral sclerosis, and epileptic patients. The results showed that the affected pathways shown by differential tractography matched well with the unique clinical symptoms of the patients, and the false discovery rate of the findings could be estimated using a sham setting to provide a reliability measurement. This novel approach enables a quantitative and objective method to monitor neuronal injury in individuals, allowing for diagnostic and prognostic evaluation of brain diseases.

## Introduction

Magnetic resonance imaging (MRI) is a commonly used neuroimaging technique for revealing a structural change in patients with neurological disorders. Studies have used structural MRI to reveal gray matter atrophy in patients with multiple sclerosis (Rovira et al., 2015; Wattjes et al., 2015) and atrophy in the caudate in patients with Huntington disease (Tabrizi et al., 2009; Tabrizi et al., 2012). In addition to structural MRI, diffusion MRI has also been explored as an imaging biomarker for early-stage neuronal injury before atrophy happens. Animal studies have used diffusion tensor imaging (DTI)(Basser et al., 1994) to detect acute demyelination or axonal loss (Song et al., 2002; Song et al., 2005). The decrease of anisotropic diffusion has been shown to be correlated with axonal loss (Budde et al., 2009; Huisman et al., 2004; Werring et al., 1999; Werring et al., 2000). However, anisotropy remains a voxel-based measurement, which is prone to local variations such as partial volume effect (Henf et al., 2018; Wang et al., 2011) or signal noise, thereby limiting its clinical applications (Melonakos et al., 2011). Greater specificity and sensitivity could be achieved by aggregating voxel-wise anisotropy changes along fiber pathways and grouping them into a “track”.

To this end, we propose a novel method called “differential tractography” to provide a track-based biomarker for neuronal injury. This method compares repeat scans of the same individuals to capture neuronal injury reflected by a decrease of anisotropic diffusion, or “anisotropy” (Fig 1a-1c). To achieve a higher specificity, we imbued the deterministic fiber tracking algorithm (Yeh et al., 2013) with a novel “*tracking-the-difference*” paradigm. This was realized by adding an additional criterion to track along trajectories on which a decrease of anisotropy was found between repeat scans (Fig. 1d-1e). Integrating this “tracking-the-difference” paradigm into the fiber tracking process resulted in a new tractography modality that tracks the exact portion of pathways exhibiting substantial differences in anisotropy. The additional criterion ignored unaffected regions and enhanced meaningful findings related to neuronal injury. In comparison, the conventional fiber tracking (Wedeen et al., 2012) is based on a “*tracking-the-existence*” paradigm. It only considers anisotropy from one MRI scan and thus will include all existing pathways regardless of whether they have an injury.

**Fig. 1.**
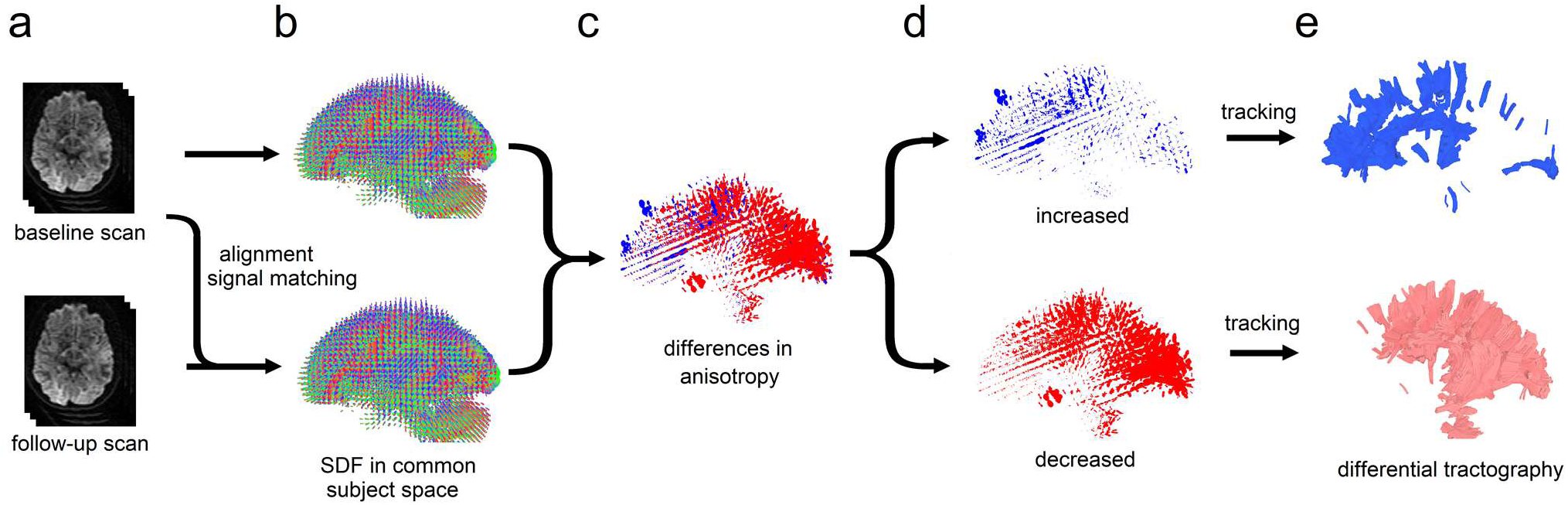
The flow chart of differential tractography. (a) The baseline and follow-up scans of the same subject are spatially aligned, and the diffusion signals are scaled to the same unit. (b) The spin distribution function (SDF) from two scans are reconstructed in the same common subject space. (c) The difference in the anisotropic component of SDF is computed for each fiber orientation. (d) Increased and decreased anisotropy values are separated to guide a “tracking-the-difference” algorithm. (e) Differential tractography shows the exact segment of tracks with increased and decreased anisotropy, respectively. The tracks with decreased anisotropy suggest possible neuronal injury, whereas the number of tracks with increased anisotropy can be used to estimate the number of false findings.

To implement differential tractography, the algorithm requires one anisotropy value for each fiber population to calculate its longitudinal change, but the fractional anisotropy (FA) derived from DTI is a voxel-based measurement, and thus all fiber orientations within the same voxel will inherit the same anisotropy value. To overcome this limitation, we used the anisotropic component of the spin distribution function (SDF)(Yeh et al., 2010) as an anisotropy measurement for each fiber population. SDF could provide one anisotropy measurement for each fiber population. This approach has been shown to be more robust against partial volume effect (Yeh et al., 2013) and achieved high accuracy in a recent competition study (ID#3)(Maier-Hein et al., 2017).

To further maximize the detection power, we used a diffusion MRI acquisition that sampled 22 diffusion sensitizations (b-values) at 257 directions, a substantial improvement over one sensitization at 30~60 directions used in the conventional settings, or 3 diffusion sensitizations at 180 directions used in the current mainstream studies (Glasser et al., 2016). The higher number of diffusion sensitizations greatly increased the chance to detect neuronal injury that involves only a subtle change in the restricted diffusion (Wang et al., 2011). We also introduce a novel sham setting that can estimate the false discovery rate (FDR) of differential tractography to provide a reliability measurement against local random error.

To evaluate the performance, we applied differential tractography to patients with four different clinical scenarios at different stages of neuronal injury (demographic information listed in Table 1). The first scenario was multiple sclerosis (MS) with the first episode of optic neuritis. The baseline scans were acquired right after the onset of the visual symptom, and the follow-up diffusion MRI scans were acquired 6 months after. This scenario tested differential tractography at the early stage of neuronal injury to explore its sensitivity, and any meaningful findings should be located near the visual pathways. The second scenario was the manifested Huntington disease (HD) with worsening clinical motor scores during the interval of their repeat MRI scans. We examined whether differential tractography could detect progressing neuronal injury at striatal pathways that are commonly affected by the disease. The third scenario studied the neuronal injury in an amyotrophic lateral sclerosis (ALS) patient with a deteriorating functional motor score. We examined whether differential tractography could be correlated with the patient’s clinical presentation. Last, we applied differential tractography to an epileptic patient treated by anterior temporal lobectomy. The baseline scan was acquired before the surgery, and the follow-up scan was acquired one year after the surgery. This examined whether differential tractography could correctly locate pathways with established neuronal injury after surgery, and meaningful findings should be in pathways previously connected to the area of resection. We also applied differential tractography to a normal subject to demonstrate how differential tractography may capture false results.

**Table 1:** Patient Demographics and Major Symptoms.

|  | diagnosis | age | sex | MRI scans | Major symptoms |
| --- | --- | --- | --- | --- | --- |
| #1 | multiple sclerosis | 44 | F | onset of symptom and 6-month follow-up | acute onset of left ocular pain, pain with eye movements, blurring vision of the left eye (20/400), loss of visual field in all quadrants. |
| #2 | multiple sclerosis | 24 | F | onset of symptom and 6-month follow-up | acute onset of left ocular pain, pain with eye movements, blurring vision of the right eye (20/125), superior altitudinal visual field defect. |
| #3 | Huntington disease | 60 | M | two scans at 5 months apart during the manifest stage | body bradykinesia on both sides<br>UHDRS motor: 45→49 (worsening) |
| #4 | Huntington disease | 55 | F | two scans at 5 months apart during the manifest stage | dystonia of left upper extremity, dragging right foot with ambulation, dystonia of the right leg<br>UHDRS motor: 53→64 (worsening) |
| #5 | ALS | 48 | M | baseline scan acquired 30 months after onset<br>follow-up scan acquired one year after. | left-hand weakness with fasciculations<br>ALSFRS-R: 45→32 (worsening) |
| #6 | epilepsy | 51 | M | before anterior temporal lobectomy and one-year follow-up after surgery |  |

**ALS:** amyotrophic lateral sclerosis, **UHDRS:** Unified Huntington's Disease Rating Scale, **ALSFRS-R:** Amyotrophic Lateral Sclerosis Functional Rating Scale-Revised

## Material and Methods

### MRI experiments on clinical patients with neurological disorders

The diffusion MRI acquisition included a baseline scan and another follow-up scan (acquired months later) of the same subject. We acquired scans on 6 patients with different neurological diseases including MS, HD, ALS, and epilepsy, in addition to one healthy volunteer. Table 1 summarizes their basic demographic and scan interval information, and brief medical history of these patients are summarized in Supplementary Materials. The ALS patient was previously reported (Abhinav et al., 2014). The diffusion data were acquired on a 3T Tim Trio System (Siemens, Erlangen, Germany) using a pulsed-gradient spin-echo 2D echo-planar imaging sequence. A 32-channel coil was used with a head stabilizer to limit head motion. Each diffusion MRI scan acquired 22 b-values ranging from 0 to 7,000 s/mm^2^ at a total of 257 diffusion sampling direction using a q-space imaging scheme (Callaghan, 1991). The in-plane resolution and slice thickness were 2.4 mm. TE=154 ms, and TR=9500 ms. The total scanning time was 45 minutes.

This is a retrospective study, and the multi-band sequence was not available previously. Currently, the same protocol using multi-band sequence has a much shorter scanning time of 12 minutes (see Discussion section for details).

### Quality control of diffusion MRI data

We applied a series of quality control to minimize possible false results due to acquisition issues. The first quality control was done by checking whether the image dimension, resolution, and b-table were consistent between repeat scans. All scan data were confirmed to have a consistent setting between repeat scans.

The second quality control was done by calculating the mean Pearson correlation coefficient of the “neighboring” diffusion-weighted images:

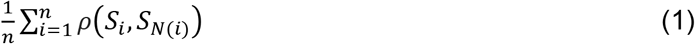

where *ρ* calculates the Pearson correlation coefficient, *S*_*i*_ is the *i*-th diffusion weighted image, and *N*(*i*) returns the index of neigthe hboring diffusion-weighted image acquired by the most similar diffusion sensitization in the q-space:

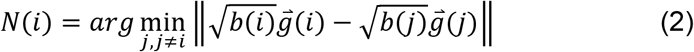

*b*(*i*) is the b-value, and 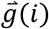 is the diffusion encoding direction. The resulting values in Eq. (1) ranges between 0.6~0.8, and a data set would be rejected if the baseline and follow-up scans have a difference in mean Pearson correlation coefficient greater than 0.1. In this study, one ALS patient’s data we recruited previously (Abhinav et al., 2014) was rejected because the baseline scan had a substantially lower value due to slice shift during the scan.

The third quality control step was identifying slice-wise signal dropout for each slice in each diffusion-weighted image. This examination excluded slices with a large background region greater than 15/16 of the entire slice area (defined by the brain mask). For each slice, we calculated its Pearson correlation coefficients with four of its signal-related slices, including its upper and lower adjacent slices and the same-location slices of two neighboring diffusion-weighted images defined by Eq. (2). The maximum of these four correlation values was used as the representative correlation coefficient of the slice, and a signal dropout would result in a decrease of this representative correlation coefficient. If a slice had an average decrease of representative correlation coefficient greater than 0.1 in comparison with its four related slices, we identified it as a signal dropout slice. In this study, we accepted the data set if the number of signal dropout slices was smaller than 0.1% of the total slice number (i.e. < 25 slices). All data set were screened and passed this criterion.

The fourth quality control was checking the b-table orientation using the fiber coherence index (Schilling et al., 2019). One scan was found to have slices order flipped upside down. The slice order and b-table were corrected before further analysis. The above-mentioned quality control routines are available on DSI Studio (http://dsi-studio.labsolver.org).

After the quality control steps, the diffusion data of the follow-up scans were compared with baseline scans using the following analysis:

### Empirical distribution of water diffusion

The empirical distribution of water diffusion was calculated from diffusion-weighted signals using generalized q-sampling imaging (GQI)(Yeh et al., 2010). This “empirical distribution” has no assumption of the underlying distribution (e.g. Gaussian distribution), and thus it can be applied to a variety of fiber or biological conditions. The empirical distribution calculated from GQI, termed spin distribution function (SDF), has a different physical definition from the *diffusivity* calculated from DTI that quantifies how fast the diffusion is. SDF quantifies the accumulated spin density of restricted diffusion sampled at any orientations, and it can be calculated using the formula:

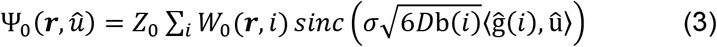

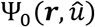 is the SDF value oriented at 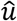 and sampled from a voxel located at ***r***. *Z*_0_ is a scaling constant to convert the arbitrary unit of the diffusion signals to a density unit. *i* iterates through each diffusion weighted signals W(***r***, i), and b(i) is the b-value, 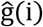 is the direction of the diffusion sensitization gradient. σ is the diffusion sampling ratio controlling the displacement range of the diffusing spins (1.25 was used in this study). *D* is the diffusivity of free water at room temperature.

We then calculated the SDFs of the follow-up scan and transformed them into the space of the baseline scan (Fig. 1a and Fig. 1b) so that they could be directly compared. This was done using q-space diffeomorphic reconstruction (QSDR)(Yeh and Tseng, 2011), a method that generalized GQI to accept spatial transformation in the reconstruction. QSDR allowed us to simultaneously reconstruct and transform SDF from the follow-up scan to the space of the baseline scan using the following formula:

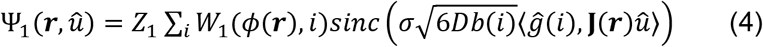

where *ϕ*(***r***) transforms spatial coordinate ***r*** from the space of the baseline scan to that of follow-up scan. *W*_1_(*ϕ*(***r***), *i*) is the diffusion-weighted signals at coordinate *ϕ*(***r***). ***J***(***r***) is the Jacobian matrix at the same coordinate that rotates the unit vector 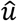. There other variables follow the same notations in Eq. (3).

Since the scans were from the same subject, we assumed there was only “rigid body” transformation (i.e. only rotation or translocation) between the scans, and the transformation was a simple matrix-vector multiplication. Please note that this assumption could be violated if there was a large tissue distortion due to edema or tissue removal, and a nonlinear spatial registration should be used in QSDR to handle this problem. The rigid body transformation matrix was obtained by linear registering the b0 images (or the sum of all diffusion-weighted images). We used the negative of the correlation coefficient between the images from baseline and follow-up scans as a cost function to calculate the transformation matrix. The cost function was minimized using a gradient descent method. The rotation matrix of the rigid body transformation was used as the Jacobian for Eq. (4).

Please note that the SDFs calculated from Eq. (3) and (4) have “arbitrary units”. Therefore, the *Z*_1_ constant in Eq. (4) had to be scaled to match the same unit of *Z*_0_ in Eq. (3). This signal matching was done using the sum of all diffusion-weighted images from two scans:

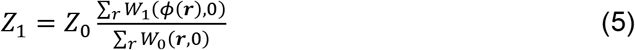

The isotropic component of an SDF was then removed by subtracting its minimum values.

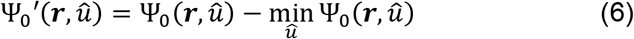

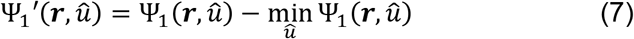

Equation (6) and (7) provide the anisotropic component of SDF (termed anisotropy hereafter) to minimize the effect of free water diffusion (Yeh et al., 2013). It is noteworthy that this anisotropy measurement has a different physical meaning from the fractional anisotropy (FA) calculated in DTI. FA is a *ratio* between 0 and 1 calculated from diffusivities and has no unit. The anisotropic SDF has the same physical unit of the SDF, which is the spin density of diffusing water.

### Tracking differences in the SDF

To track differences along the existing fiber pathways, we first determined the local fiber orientations using the peaks on the sum of 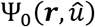 and 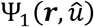, and then the anisotropy estimated from the summed SDF was used to filter out noisy fibers and to define the termination of the white matter tracks, as what had been done in conventional tractography (Yeh et al., 2013). The percentage difference in the anisotropy between baseline and follow-up scans was then calculated (Fig. 1c):

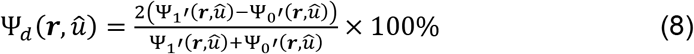

The percentage changes in the anisotropy, 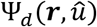 can have positive values (blue SDFs in Fig. 1c), which indicates an increase in the density of anisotropic diffusion, or negative values (red SDFs in Fig. 1c), which indicates a decrease in the density of anisotropic diffusion.

An additional tracking-the-differences criterion was added to the fiber tracking algorithm to track the exact segment with a decrease or an increase in the anisotropy greater than a *change* threshold, Specifically, to track pathways with an increase of anisotropy, the additional criterion checked whether the increase of anisotropy was greater than a predefined value of percentage change (e.g. 20%) and continued tracking as long as the criterion was satisfied:

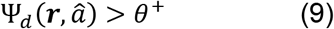

where 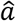 is the local fiber directions used in the fiber tracking algorithm. Similarly, to track pathways with decreased anisotropy, the criteria continued tracking if the decrease of anisotropy was greater than a predefined value of percentage change (e.g. −20%):

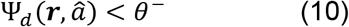

This allowed us to track two different sets of pathways, one for increased anisotropy, one for decreased anisotropy. The other existing criteria in conventional tractography (e.g. seeding strategy, propagation interval, angular threshold, length constraint…etc.) remained in effect as what has been used in the generalized deterministic fiber tracking algorithm (Yeh et al., 2013). It is noteworthy that the angular and anisotropy thresholds in the original tracking algorithm were still used in differential tractography to eliminate noisy fiber and to ensure a correct white matter coverage. The criteria (9) or (10) (termed “positive change threshold” and “negative change threshold”) served as *additional* constraints to limit the findings to the exact segment of pathways with a substantial change in the anisotropy value.

In this study, the differential tractogram was obtained by placing a total of 5,000,000 seeding points in the white matter. The angular threshold was randomly selected between 15 to 90 degrees. The step size was 1 mm, and the anisotropy threshold was automatically determined by DSI Studio. Two iterations of topology-informed pruning (Yeh et al., 2018) were applied to the tractography to remove noisy findings. The above-mentioned setting was regularly used in conventional tractography. We tested differential tractography with different values of change threshold (5%, 10%, 15% …. 50%) and length threshold (5 mm, 10 mm, 15 mm, … 50 mm). Track with lengths shorter than the length threshold was discarded, and the results of different length threshold and change threshold were compared to access its effect on the sensitivity and specificity of differential tractography.

### Estimating false discovery rate using a sham setting

Figure 2 illustrates the experimental design that allows for estimating the false discovery rate (FDR) of differential tractography in individual scans. As shown in Fig. 2a, the baseline scan is compared with the follow-up scan (upper row) using differential tractography to reveal tracks with decreased anisotropy (Fig. 2b, upper row). If a total of 5,000,000 tracking iterations are conducted, we can view each of them as an independent hypothesis testing since each tracking iteration is done independently. The null hypothesis is “there exists no track with a decrease in the anisotropy”. This null hypothesis will be rejected if the track length is longer than a predefined length threshold (e.g. 40 mm). Each rejected hypothesis is thus regarded as a “positive finding”, but here the finding can be either a true positive or false positive. As shown by the example in the upper row of Fig. 2b, the number of tracks with a length longer than 40 mm is 13947, meaning that there are 13947 rejected hypotheses as positive findings. These 13947 findings include true positive and false positive, and we will need to estimate the number of false-positive findings so that FDR can be estimated. To estimate the number of false-positive findings, a sham scan can be acquired on the same day of the baseline scan or within a timeframe that no neuronal change is expected (Fig. 2a, lower row), and thus any positive findings shown in the sham scan should be false-positive findings. The sham scan is also compared with the baseline scan, and the same number of 5,000,000 tracking iterations are conducted to see how many positive findings are generated. The example in the lower row of Fig. 2b shows that the sham scan generates a total of 177 tracks with a length longer than 40 mm, meaning that the estimated number of false-positive findings is 177. Using this information, we can calculate the FDR, which is 177/13947=0.0127 (Fig. 2c).

**Fig. 2.**
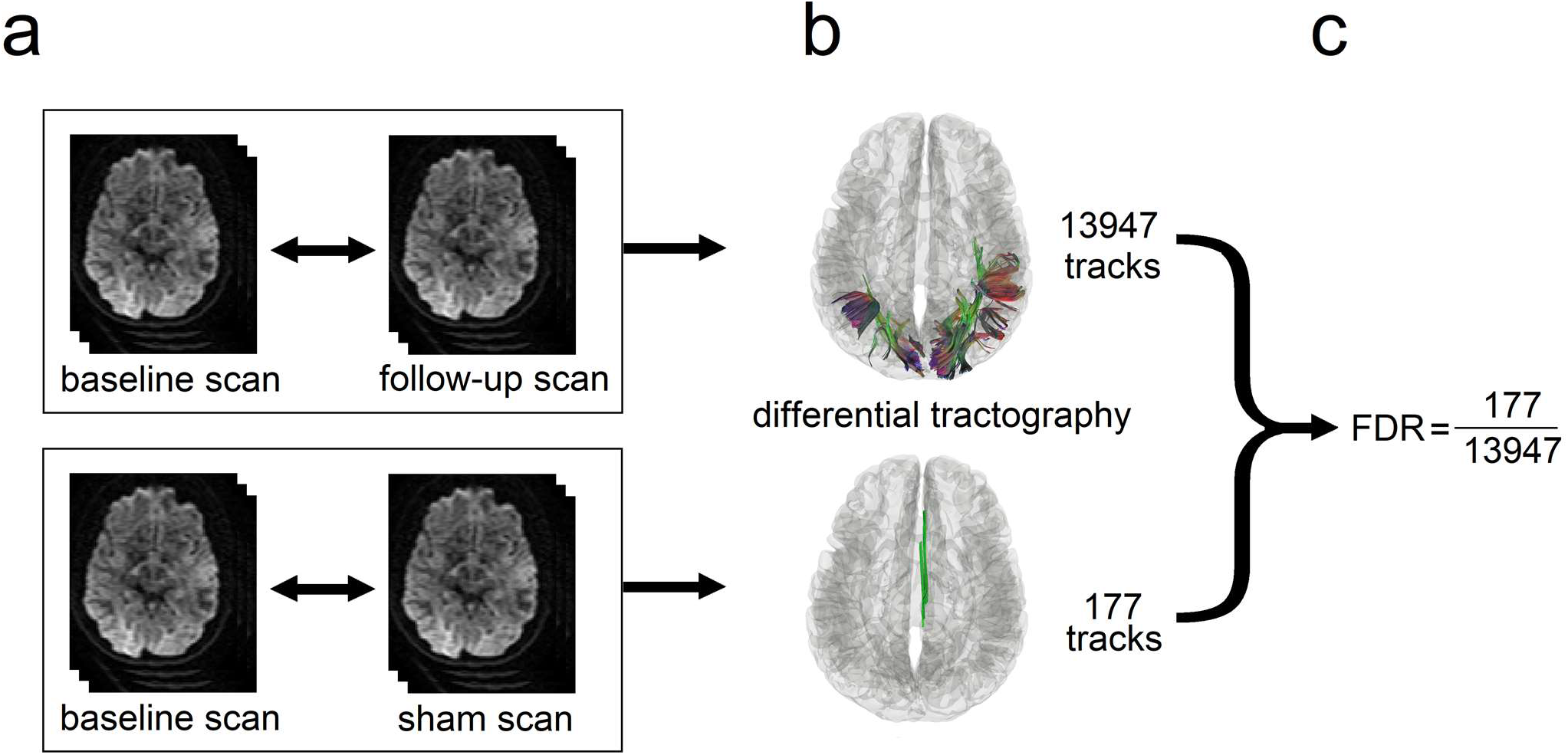
Diagram showing the sham setting for calculating the false discovery rate (FDR). (a) The baseline scan is compared with a follow-up scan (upper row) and a sham scan (lower row), respectively. The follow-up scan is often acquired months after the baseline scan to capture the true positive findings, whereas the sham scan can be a repeat scan on the same day or any scan setting that ensures the findings are all false positives. (b) The number of findings from the follow-up scan (upper row) includes both true and false positives, whereas the number from the sham scan (lower row) include only false-positive findings. (c) These two numbers can be used to estimate FDR.

### Substitute sham

In this retrospective study, we did not have an additional sham scan, and a “substitute sham” approach was used. We assumed that there should be no increased track integrity during the disease course, any findings showing *increased* anisotropy can be regarded as false-positive findings. Therefore, we can use the number of tracks with *increased* anisotropy in the follow-up scan as a substitute for the sham scan. Even if tracks with increased integrity do exist due to re-myelination, it will increase our estimated number of false-positive findings. This means that the FDR estimated by this substitute sham approach will be an upper bound of the actual FDR. Therefore, instead of reporting “FDR=0.0127”, the FDR calculated using a substitute sham should be reported as “FDR ≤ 0.0127”.

The processing pipeline for differential tractography and the quality control procedure is implemented in DSI Studio (http://dsi-studio.labsolver.org). Documentation and source code to reproduce the same result is also available on the website.

## Results

### Neuronal injury reflected by a decrease of anisotropy

Figure 3a shows the intermediate results of differential tractography applied to an MS patient with optic neuritis (patient #1, demographics summarized in Table 1). The baseline scan was acquired right after the onset, whereas the follow-up scan was acquired 6 months after. For each fiber orientation in a voxel, differential tractography compares the anisotropy differences between two MRI scans in a common subject space (Fig. 1a-1c). The fiber orientations with a decrease of anisotropy greater than 30% could are plotted by red sticks in Fig. 3a. In the figure, most of the differences are distributed near the primary visual pathways, whereas some spurious differences are randomly distributed throughout the entire whiter matter regions, most likely due to local signal variations or registration error.

**Fig. 3.**
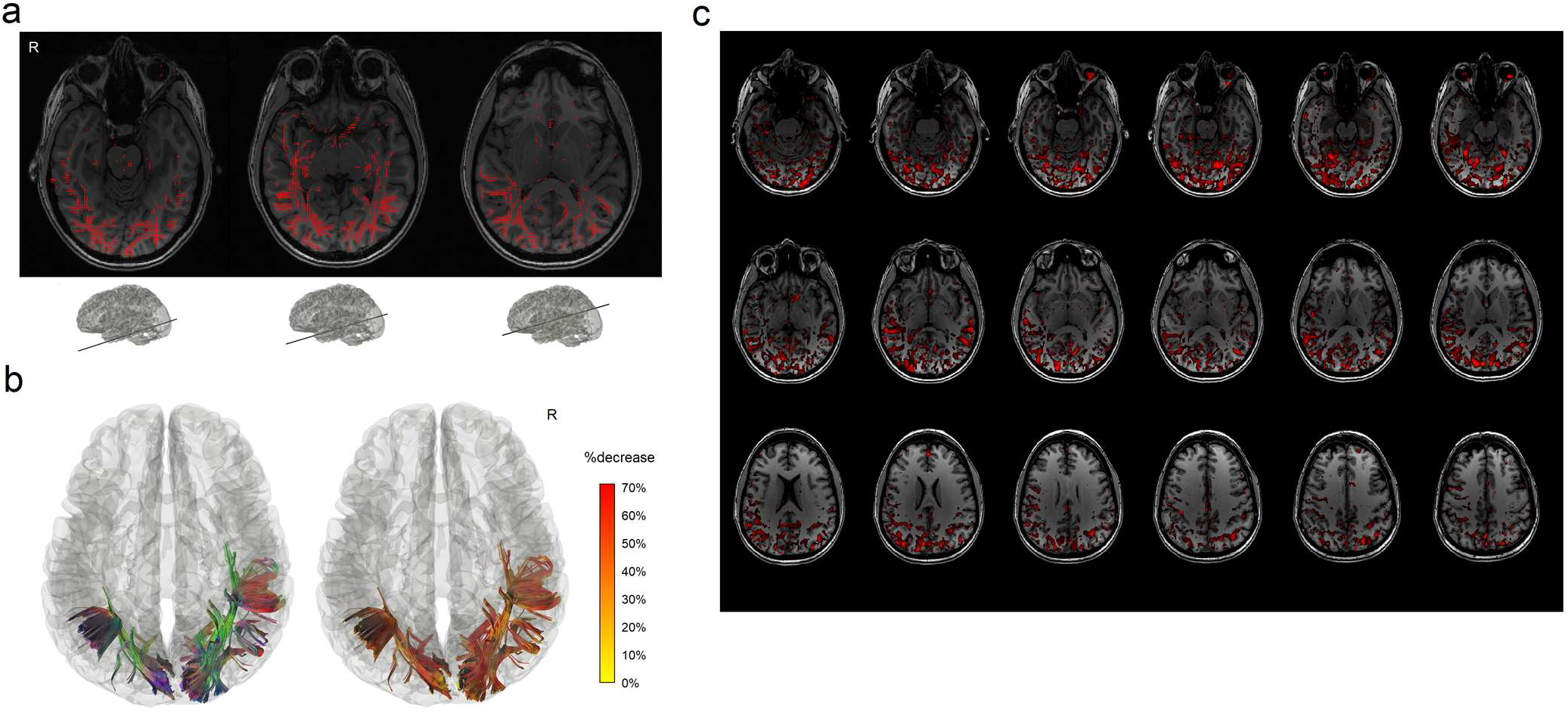
Differential tractography of a multiple sclerosis patient with the first episode of optic neuritis. (a) The intermediate result of differential tractography shows red sticks indicating local fiber orientations with a negative change threshold of 30% between repeat scans. The sticks are mostly distributed along the primary visual pathways, while sporadic false findings can also be found throughout the entire whiter matter regions due to local signal variations. (b) The red sticks are tracked and connected into continuous trajectories, whereas the other unaffected parts of the white matter pathways are ignored. The resulting 3D presentation is the differential tractogram of the patient showing the exact segment of pathways with a substantial decrease in anisotropy. The tractography can be rendered by directional colors (left) or severity-coded color (right) to provide information about the spatial location, and the severity of the axonal damage can be quantified by percentage decrease of anisotropic diffusion. (c) The same data analyzed by voxel-based differences show numerous fragmented findings possibly due to numerous local random error. There is no track information to assist correlating structure with a function and differentiating true findings from false ones.

To eliminate these local spurious differences, the “tracking-the-difference” algorithm is applied to the track and link all local differences together into continuous trajectories, and short fragments are discarded using a length threshold of 40 mm (Fig. 3b). The rationale behind this length threshold is that the local random error does not propagate along fiber pathways, whereas true findings due to neuronal injury will form a continuous decrease of anisotropy along the fiber bundles. A length threshold will effectively differentiate between them and help eliminate false results.

The left inset figure in Fig. 3b shows affected tracks in directional colors (red: left-right green: anterior-posterior blue: superior-inferior), whereas the tracks in the right inset figure are color-coded by the percentage decrease of anisotropy suggesting the severity of neuronal injury (yellow: 0% decrease red: 70% decrease). In overall, the differential tractogram in Fig. 3b reveals a heterogeneous decrease of anisotropy between 20% to 50%. All findings are in the bilateral primary visual pathways or their collateral connections. The location of the finding matches well with the patient’s medical history of visual field loss in both left and right quadrants. The topology of affected pathways seems to present a ripple effect: not only the primary visual pathway is affected, but also certain collateral connections to the visual cortex has shown a decrease in the anisotropy. Although this patient was fully recovered from the symptoms during the follow-up scan (brief medical history in the Supplementary materials), differential tractography still captures subclinical change near the bilateral optic radiation.

We further compare differential tractography with conventional voxel-wise statistics. Figure 3c shows the axial mapping of anisotropy differences for each voxel using the same data. The red regions are voxels with anisotropy decrease greater than 30%, and the numerous fragments can be observed across the entire brain regions. Those fragments could be due to local random error and thus may not be true findings. This illustrates a typical limitation of a voxel-based imaging biomarker, and there is no pathway information to assist diagnostic evaluation. In comparison, the differential tractogram in Fig. 3b provides track-based imaging biomarkers that can easily assist diagnostic evaluation by offering trajectories information. The findings can be associated with an anatomical pathway to infer the functional correlation.

### Conventional tractography versus differential tractography

Figure 4 compares conventional tractography (Fig. 4a) with differential tractography (Fig. 4b) on another MS patient with optic neuritis (patient #2 demographics summarized in Table 1). The conventional tractography was generated using the baseline scan, whereas the differential tractography was configured to map pathways with more than a 30% decrease in anisotropy with a length threshold of 40 mm. The trajectories in Fig. 4a and Fig. 4b are both colored-coded with directional colors (red: left-right green: anterior-posterior blue: superior-inferior). The first row shows tractography viewing from the top, whereas the second rows show from the left. The conventional tractography in Fig. 4a visualizes the trajectories of the entire brain fiber pathways, and there is no gross anomaly visible from the tractography that may suggest a major neurological disorder. This is expected because, at the early stage of multiple sclerosis, the patient usually does not present a gross structural change that can be readily identified in conventional tractography. In comparison, the different tractography in Fig. 4b pinpoints the location of affected pathways in the bilateral primary visual pathway near the visual cortex. The location matches well with the patient’s disease presentation of optic neuritis, whereas conventional tractography in Fig. 4a shows no clue to this critical information.

**Fig. 4.**
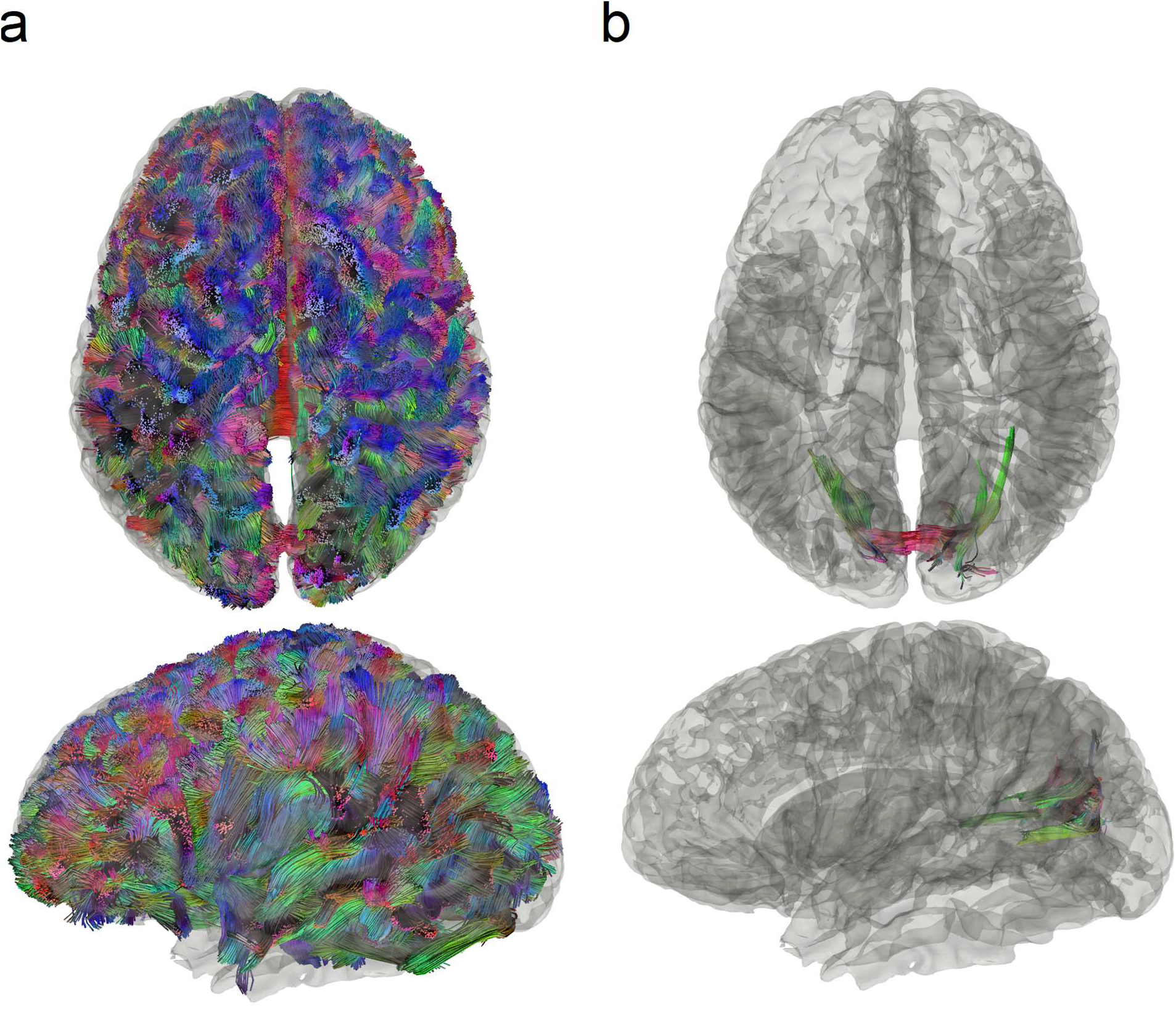
Conventional tractography compared with differential tractography on a multiple sclerosis patient with the first episode of optic neuritis. (a) Conventional tractography shows all existing fiber pathways in the human brain and is insensitive to any subtle decrease in the diffusion property. (b) Differential tractography ignores unaffected regions and shows the exact segments of the pathways that have a substantial decrease of anisotropy quantified between repeat scans of the same individual.

### Reliability assessment

We further use patient #1 and a 42-year-old normal subject as the examples to show how the reliability of differential tractography findings can be quantified using a sham setting. Both the follow-up scans of these two subjects were acquired 6 months after the baseline scans. Figure 5a is the differential tractogram of patient #1 showing pathways with an increase or decrease anisotropy greater than 30% at different length thresholds, whereas Fig. 5b is the same analysis applied to a healthy subject. Only the decreased anisotropy in patient #1 (lower row in Fig. 5a) contains possible true findings for neuronal injury, whereas other rows in Fig. 5a and Fig. 5b are all false-positive results due to either physiological noises (cardiovascular or respiratory) or phase distortion artifact. Most false findings can be effectively removed using a longer length threshold, and there is a trade-off between sensitivity and specificity controlled by the length threshold. A longer length threshold renders a more specific result with the expense of losing meaningful findings in shorter segments, whereas a shorter length threshold allows for more findings with a risk of taking false results. To quantify the reliability of findings in the lower row of Fig. 5a, we used the total number of findings at the upper row of Fig. 5a as an estimation for the number of false findings. This assumes that the random error will produce a similar number of false findings in both rows and allows us to estimate FDR.

**Fig. 5.**
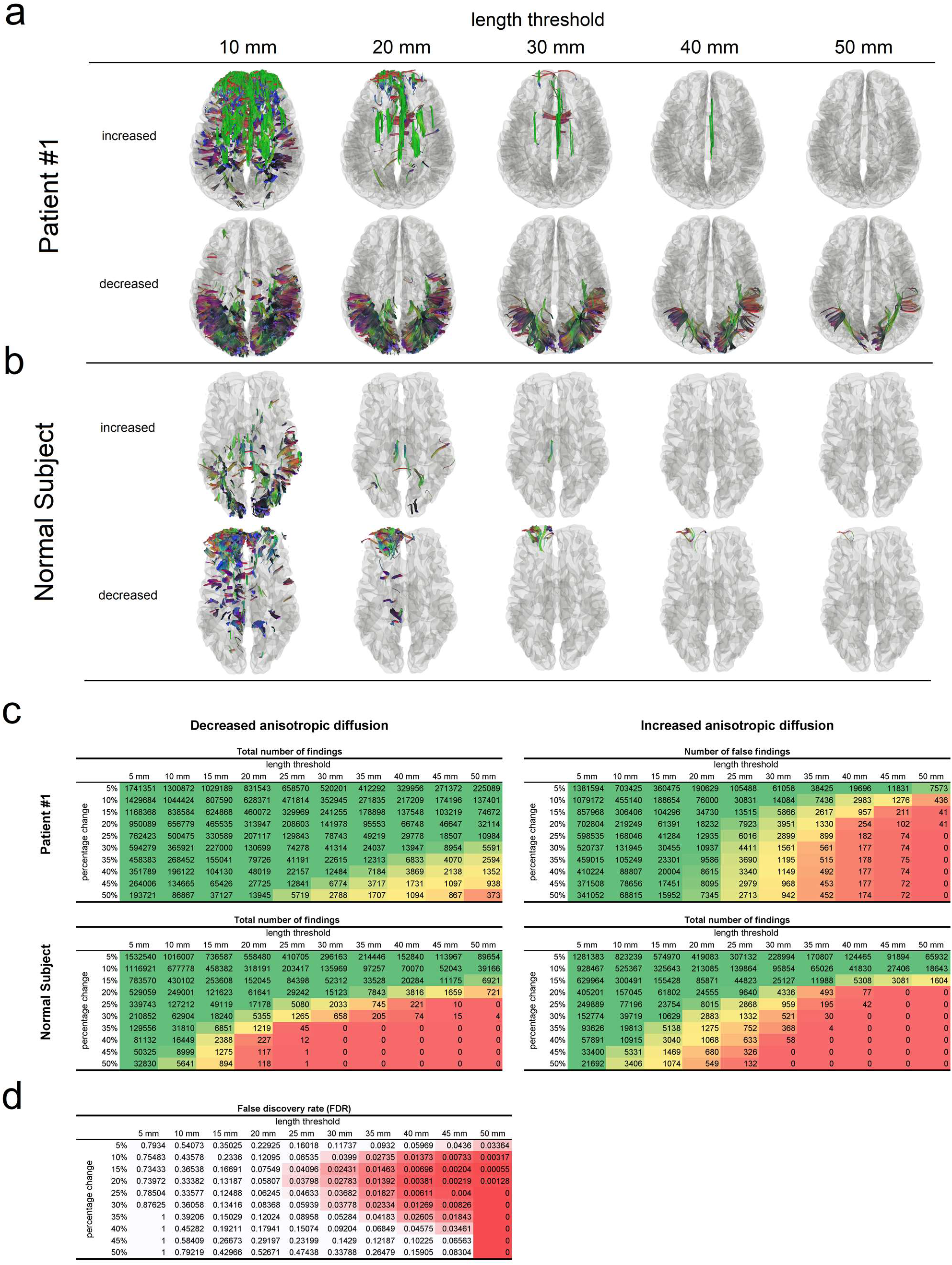
Reliability assessment of differential tractography using the length threshold. (a) Differential tractography is applied to a multiple sclerosis patient at different length thresholds. Only the tracks with decreased anisotropy in the patient may contain true positive findings. A longer length threshold (e.g. > 40mm) can reduce false findings at the expense of sensitivity, whereas a shorter threshold may introduce more false results. (b) Differential tractography is applied to a normal subject, and any findings in the normal subject are false positives for neuronal injury. (c) The numbers of findings at different length thresholds and change thresholds are listed in tables. The patient has substantially large numbers of tracks with decreased anisotropy, suggesting a possible neurological injury. In comparison, the normal subject has similar numbers of tracks with increased and decreased anisotropy. (d) False discovery rate (FDR) of the findings in a patient can be calculated by using the patient’s own numbers of tracks with increased anisotropy as an estimation of the number of false findings. This allows for adjusting the sensitivity and specificity of differential tractography and quantifying the reliability at different length and change thresholds.

Figure 5c and 5d illustrate the FDR calculation process. Figure 5c lists the number of findings at different length thresholds and the change thresholds for patient #1 (upper two tables) and the normal subject (lower two tables). For example, a negative change threshold of 30% means that the tracking algorithm will track only the fiber orientations with a decrease of anisotropy greater than 30%. The tables on the left are numbers of findings with decreased anisotropy, whereas those on the right are numbers of findings with increased anisotropy. The green colors in the table are those with a larger number of findings, whereas the red colors indicate a smaller number. The tables of the patient show a substantially larger number of findings with decreased anisotropy caused by the disease. In comparison, the two tables of the control subject are substantially similar, and the false results are equally distributed in both tables. This result supports the usage of the “substitute sham” approach mentioned in the Material and Methods section: we can reasonably use the findings with increased anisotropy (the table on the right) of the same subject to estimate the number of false findings for calculating FDR.

Figure 5d shows the FDR of patient #1’s findings calculated using the “substitute sham” (the increase of anisotropy results from the same subject). The resulting table shows a trade-off between sensitivity and specificity controlled by both the length threshold and change threshold. A length threshold of 30~40 mm and a change threshold at 20~30% decrease of anisotropy provides us an FDR around 0.01, suggesting that around 1% of the tracks shown in the differential tractogram are false results. We can use these two thresholds to leverage sensitivity and specificity. For example, lower thresholds are geared toward higher sensitivity to explore potential neuronal injury, whereas higher thresholds can provide a confirmatory answer to the axonal damage. The optimal setting can be different based on the disease condition, scan interval, and purposes (e.g. exploratory or confirmatory).

### Differential tractography on patients with neurological diseases

We further apply differential tractography to patients with different neurological disorders in Fig. 6 and list the FDR of these findings in Table 2. The scan subjects include patients with MS (#1, #2), HD (#3, #4), ALS (#5), and epilepsy (#6). The first three rows show the differential tractograms in three views (left sagittal from left, coronal from the front and axial from top) using directional colors, where the last row shows the differential tractogram with yellow-red colors representing the percentage decrease of anisotropy.

**Table 2:** False Discovery Rate of Differential Tractography Findings.

|  | diagnosis | change<br>threshold | length<br>threshold | number of<br>findings | false-positive<br>findings* | FDR |
| --- | --- | --- | --- | --- | --- | --- |
| #1 | multiple sclerosis | 30% | 40 mm | 13,947 | 177 | $\leq 0.0126$ |
| #2 | multiple sclerosis | 30% | 40 mm | 2,799 | 0 | $< 0.0001$ |
| #3 | Huntington disease | 30% | 40 mm | 85,243 | 3,571 | $\leq 0.0419$ |
| #4 | Huntington disease | 30% | 40 mm | 64,272 | 0 | $< 0.0001$ |
| #5 | ALS | 15%† | 40 mm | 12,222 | 2,548 | $\leq 0.2085$ |
| #6 | epilepsy post-op | 30% | 40 mm | 15,959 | 0 | $< 0.0001$ |
|  | normal control | 30% | 40 mm | 74 |  |  |

\* estimated by the number of tracks with increased anisotropic diffusion
†30% yields no findings

**Fig. 6.**
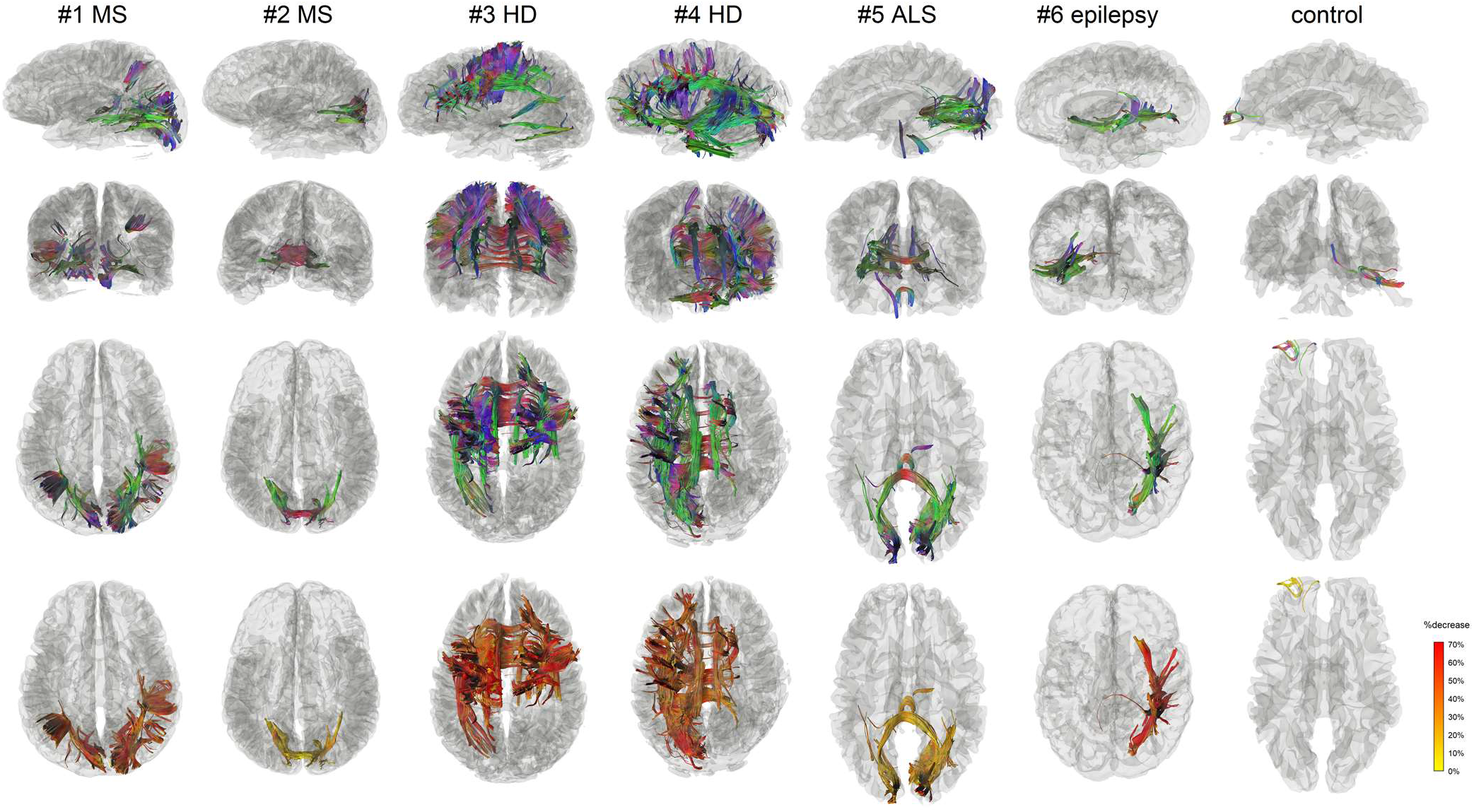
Differential tractograms of patients with different neurological disorders in comparison with a normal subject. The results were generated automatically without expert selection. The differential tractograms of the two MS patients match well with their clinical presentation of optic neuritis. Patient #1 has a much severer drop in visual acuity, which can be quantitatively reflected by her larger the volume of the findings and a greater percentage decrease of the anisotropy along the affected pathways. The differential tractograms of the two Huntington diseases show extensive affected striatal pathways. Patient #4 had more asymmetric dystonia, matching the asymmetry presentation of the differential tractography. The ALS patient had lower motor neuron presentation of left-hand weakness, matching the finding of right lower corticospinal pathways in differential tractography. The epileptic patients received right anterior temporal lobectomy, matching the findings in the differential tractogram that shows the affected pathways around the surgical location. The false findings in the normal subject can be differentiated by their less significant decrease of anisotropy and their location at the anterior frontal region, which is known to be more susceptible to phase distortion artifact.

The first notable finding comparing MS patients #1 and #2 is that the volume of affected pathways and their decrease of anisotropy reflect the severity of their clinical symptoms. The medical history of patient #1 (Supplementary Materials) indicates a more severe drop in visual acuity to 20/400 in addition to her visual field defect in all quadrants, while patient #2 only had a decrease of visual acuity to 20/125 with only superior altitudinal visual field defect (Table 1 and Supplementary Materials). The greater severity in patient #1 is reflected by a larger volume of affected pathways diffusion (patient #1: 55681.6 mm^3^, patient #2: 26124 mm^3^) and a greater decrease of anisotropy shown in the last row. This suggests that differential tractography has a quantitative potential to evaluate disease severity using either the volume of affected tracks or the decrease of their anisotropy.

The differential tractogram of HD patients #3 and #4 in Fig. 6 both shows affected pathways around the stratum. This matches well with the common understanding that striatal pathways are usually involved in Huntington’s disease. Moreover, the differential tractogram in patient #4 has a greater involvement extending to brainstem and cerebellum, suggesting a worse motor performance. This seems to match the patient’s medical history of a higher motor score of 64 (Table 1 and Supplementary Materials). The patient also had more asymmetric dystonia, matching the asymmetry presentation of the differential tractography.

Patient #5 in Fig. 6 is an ALS patient. It is noteworthy that this patient had mostly lower motor neuron symptoms (weakness), and thus might not have positive findings in the brain. The differential tractogram of this patient were obtained using a 15% negative change threshold because a threshold of 30% yielded no findings. Differential tractography reveals only a minor decrease in this patient (other cases in Fig. 6 have mostly greater than 30% decrease). This could be explained by the fact that the patient had predominately lower motor neuron symptoms affecting predominately peripheral nerves. Therefore, the findings in the central nervous system could be only subclinical. Nonetheless, as we lowered the change threshold to 15%, differential tractography showed affected pathways in the right lower corticospinal pathway (blue-purple colored), superior cerebellar peduncle, and posterior corpus callosum, as shown in Fig. 6. The right corticospinal pathway involvement seems to match the patient’s history of left side involvement, but it is noteworthy that the FDR of these findings were much higher (FDR~0.2 in Table 2), meaning that around 1/5 of the findings are false positives. The corpus callosum and occipital lobe findings could be subclinical damage and did not present any clinical symptoms (to be discussed in the Discussion section).

Patient #6 was a 51-year-old male with right anterior lobectomy. He was previously an epileptic patient with recurrent epilepsy (Supplementary Materials). The MRI scans were done before the surgery and one year after the surgery. Differential tractography accurately locates the location of the surgical resection in the mesial structures and approximately 5 cm of the anterior temporal neocortex. Moreover, it further reveals the pathways that were affected by the resection. While the surgical resection only removed part of the temporal gyri, the affected pathways involve much greater connection networks. Furthermore, the last row shows that the decrease of anisotropy is mostly greater than 50%, indicating a greater axonal loss due to the surgical removal of the brain tissue.

The last column in Fig. 6 shows the differential tractogram of a control. We applied the same settings to examine how differential tractography may capture false results. The result shows a mild decrease of anisotropy as presented by yellow tracks in the last row, a clue that the change may be a false positive result. Furthermore, there are only 74 findings located at the prefrontal cortex, and these findings are relatively insignificant compared to those of the patient population that shows thousands of findings (Table 2). Moreover, the location of the findings is known to be heavily affected by the phase distortion artifact, and the findings could be due to the different level of distortion between the repeat scans. However, it is noteworthy that the calculated FDR will be 0 in this case because there are no findings with increase anisotropy for this control subject. This result suggests that the FDR estimation still have its limitation if the number of findings is sufficiently small. The interpretation of differential tractography results still needs to consider the percentage decrease of anisotropy, the total number of findings, and possible sources of imaging artifact.

In general, the findings in Fig. 6 allows us to quickly differentiates the possible locations of neuronal injury and evaluate the severity. The affected pathways in MS, HD, and ALS patients show distinctly different topology, allowing for differential diagnosis or prognosis evaluation. Table 2 further shows how we can use the sham setting to calculate FDR to evaluate the reliability of the results.

### Performance difference after dropping high b-value acquisitions

We further investigate whether high b-value signals contribute to the detection power of differential tractography. In this study, we acquired 22 b-values ranging from 0 to 7,000 s/mm^2^, but most diffusion MRI studies would acquire b-values between 1,000 and 3,000 s/mm^2^ because higher b-values result in poorer signal-to-noise (SNR) ratio. This can be observed in Fig. 7a showing the diffusion-weighted images at different b-values from patient #1. It is thus of a practical consideration to limit the b-values under a certain maximum value. To examine whether the high b-value acquisition is necessary for differential tractography, we repeated the same differential tractography analysis but used only the signals with b-values between 0 and 3,000 s/mm^2^ (a total of 85 sampling directions). The reduced dataset is very similar to the scheme acquired in a previous study (Wang et al., 2011).

**Fig. 7.**
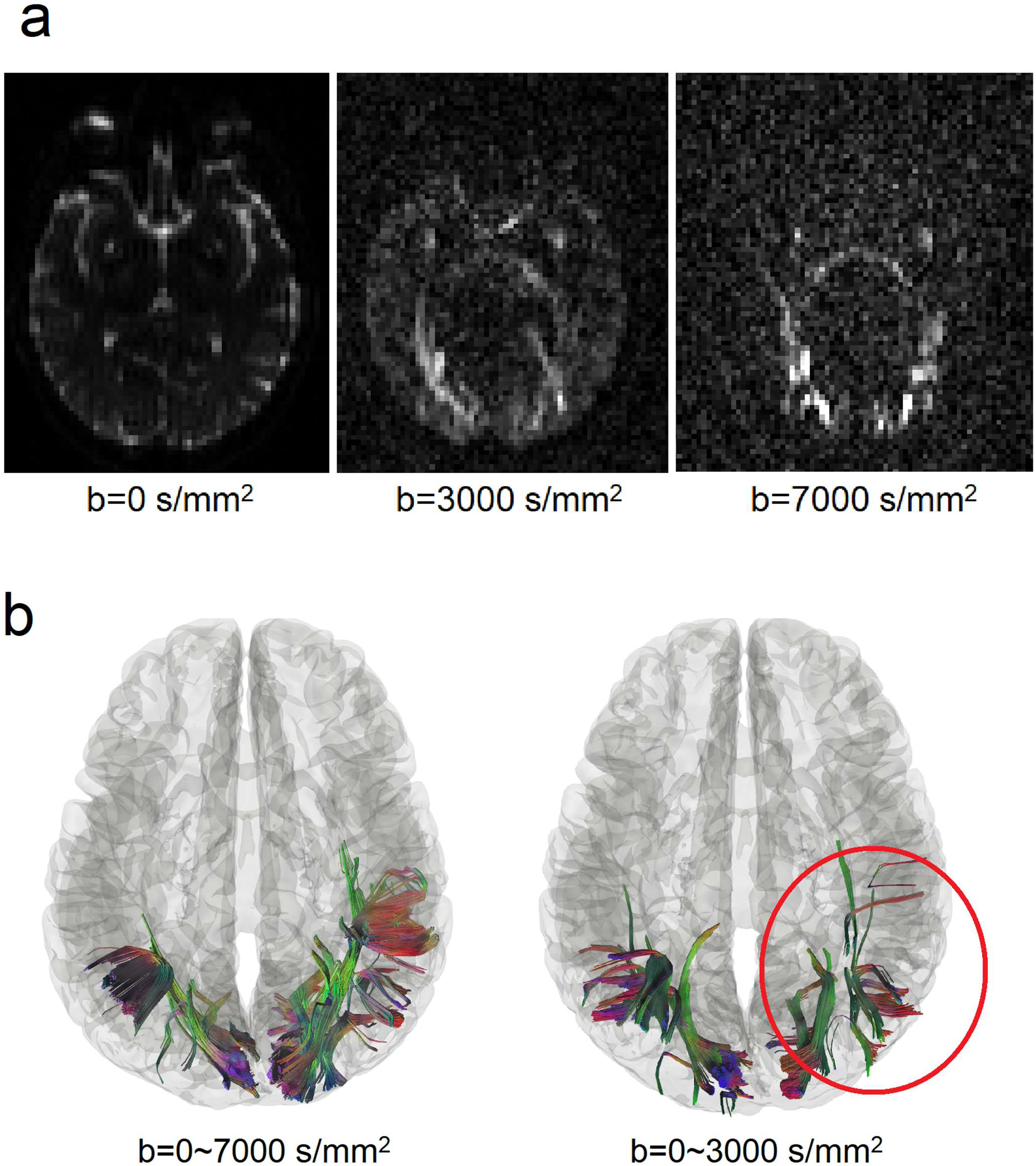
Performance differences in differential tractography due to different b-values. (a) Diffusion-weighted images of a patient at b-values of 0, 3000, and 7000 s/mm^2^. Signals at high b-values are sensitive to restricted diffusion but have a lower signal-to-noise ratio. Thus most clinical scans only acquire b-value lower than 3000 s/mm^2^. (b) Differential tractography using reduced b-values between 0 and 3,000 s/mm^2^ shows 54% fewer findings than those from the full dataset using b-values between 0 and 7,000 s/mm^2^. Although the reduced b-value dataset also shows a grossly similar result, its FDR is substantially higher (FDR=0.32) and thus not as reliable as the full dataset that includes high b-value data. This indicates the important role of high b-value acquisition in detecting early neuronal injury.

Fig. 7b shows a qualitative comparison on subject #1 before and after dropping high b-value signals. The location of the findings is mostly consistent; however, dropping high b-values results in substantial loss of the findings (annotated by the red circle). This implies that certain stages of neuronal injury can only be detected by very high b-values. The quantitative comparison is listed in Table 3. As shown in the table, there is a unanimous drop in positive findings for all patients, ranging from 38% decrease to 97% decrease, while the false findings in the normal subject increased by 7 times. Consequently, patients #1 and #3 have a dramatic FDR increase from less than 0.05 to more than 0.3, making the findings not statistically reliable. It seems that differential tractography using only low b-value signals is more sensitive to physiological variations, whereas differential tractography using both high and low b-value signals are more reliable in detecting are more specific to neuronal injury. This could be explained by the fact that the low b-value images in Fig. 7a have more signal contribution from free water diffusion in ventricles and subarachnoid space, whereas high b-value images only have signals from restricted diffusion in the core white matter. Excluding high b-value signals will shift the focus to free water diffusion and lead to more false-positive results. The overall result supports the necessity of including high b-value acquisition for differential tractography to detect neuronal injury.

**Table 3:**
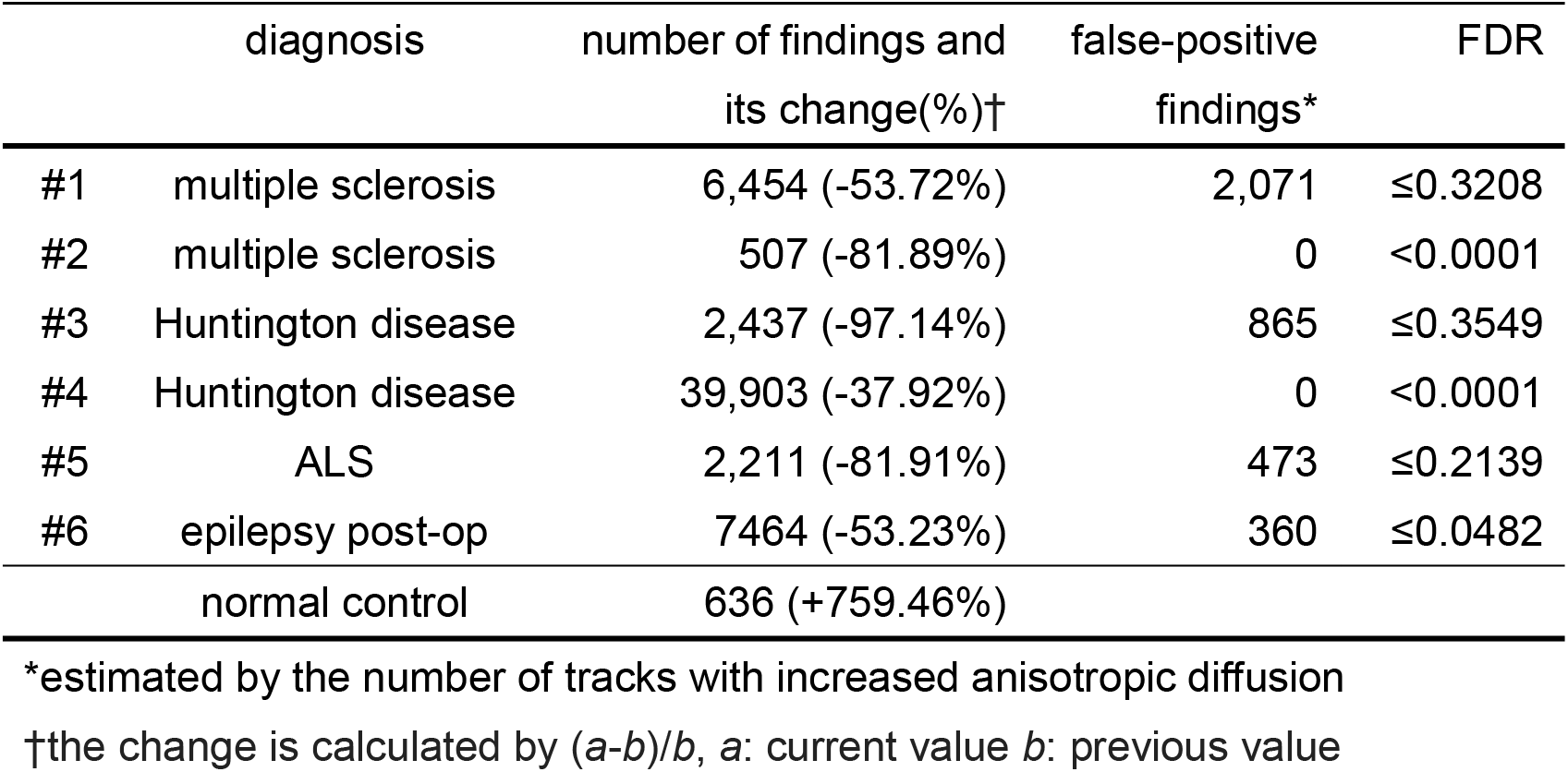
Differential Tractography Using Reduced Datasets (b = 0 ~ 3000 s/mm^2^)

|  | diagnosis | number of findings and<br>its change(%)† | false-positive<br>findings* | FDR |
| --- | --- | --- | --- | --- |
| #1 | multiple sclerosis | 6,454 (-53.72%) | 2,071 | ≤0.3208 |
| #2 | multiple sclerosis | 507 (-81.89%) | 0 | <0.0001 |
| #3 | Huntington disease | 2,437 (-97.14%) | 865 | ≤0.3549 |
| #4 | Huntington disease | 39,903 (-37.92%) | 0 | <0.0001 |
| #5 | ALS | 2,211 (-81.91%) | 473 | ≤0.2139 |
| #6 | epilepsy post-op | 7464 (-53.23%) | 360 | ≤0.0482 |
|  | normal control | 636 (+759.46%) |  |  |

\*estimated by the number of tracks with increased anisotropic diffusion
†the change is calculated by $(a-b)/b$ , $a$ : current value $b$ : previous value

## Discussion

Here we report a novel tractography method to reveal fiber pathways affected by a neuronal injury. We found that differential tractography could serve as a track-based biomarker to provide localization of neuronal injury and allow for quantifying its severity using the decrease of anisotropy and the total volume of affected pathways. The estimated FDR further offered reliability information to interpret the results, and the findings correlated well with clinical presentations of each individual.

### Comparison with other methods

Diffusion MRI fiber tracking can be viewed as a clustering process that is directionally sensitive. By using spatial relations across multiples voxels, the tracking-the-difference strategy used in differential tractography has the potential to differentiate true findings from local errors, since neuronal injury will propagate along axonal fibers while local error stays locally. This tracking-the-difference strategy is conceptually similar to clustering used in voxel-based morphometry (Ashburner and Friston, 2000) or fMRI studies (Woo et al., 2014), which groups voxel-wise statistics into clusters to achieve greater statistical power. The difference is that the clustering used in previous studies did not consider local fiber orientations and would include voxels at all possible neighboring directions, whereas differential tractography only allows findings along the fiber pathways. This improvement is similar to those using fiber geometry to increase specificity (Raffelt et al., 2017; Zhang et al., 2018), and It is likely that this structural restriction may achieve a better specificity than a regular clustering approach.

### Optimal b-table for differential tractography

The diffusion MRI acquisitions played a critical role to boost the sensitivity of differential tractography. Our result shows that acquisitions using only b-values lower than 3,000 s/mm^2^ may have a limited detection power for early neuronal injury. We also tested differential tractography on existing DTI data, and the preliminary result (not reported here) also showed a substantially higher rate of false findings. This is not surprising because a typical DTI acquisition only acquires only one b-value of 1,000~2,000 s/mm^2^ at 30~60 directions, whereas in this study, we acquired 22 b-values from 0 to 7,000 s/mm^2^ at 257 directions. It is likely that early axonal injury affects mostly restricted diffusion and can only be reliably captured if a wider range of b-value is acquired with enough diffusion sampling directions.

Multishell-acquisition could also be used by differential tractography to detect neuronal injury, but its challenges are how to ensure a homogeneous sampling density in the q-space acquisition to make the acquisition “rotation invariant”. In addition, most multi-shell acquisitions have an over-sampling problem within each shell and under-sampling problem between shells, which can be observed by plotting the sampling points in the q-space. Consequently, the inter-shell coverage may not be sufficient enough to capture a variety of diffusion patterns, while the intra-shell signals can be easily interpolated due to oversampling, meaning that there is redundancy which can be further reduced to save scanning time.

The q-space grid sampling used in this study seems to the method of choice for differential tractography because it has uniform sampling density in the q-space and covers 22 different b-values. This maximized the chance to detect a variety of diffusion pattern that could be altered during the disease process. Q-space imaging used to be criticized for its lengthy scanning time; however, after the introduction of the multi-band sequences, the updated q-space grid sampling scheme could be acquired within 12 minutes for 256 directions and 6 minutes for 128 directions, making it highly feasible for clinical studies. The exact steps to reproduce these two q-space grid acquisitions are documented on the DSI Studio website (http://dsi-studio.labsolver.org/).

### Anisotropy measurement for differential tractography

The anisotropy used in this study is not the commonly used FA provided by DTI. FA is a voxel-wise measurement, and it does not selectively quantify the anisotropy at different fiber population within the same voxel. In comparison, the anisotropy in this study can quantify the anisotropy for each fiber population at different directions This results in greater specificity to individual’s connectivity patterns (Yeh et al., 2016) and better performance in handling the partial volume effect (Yeh et al., 2013).

### Subclinical findings

Another interesting result we observed in this study is that differential tractography seems to capture subclinical findings: certain pathways could be injured without any obvious clinical symptoms reported by patients. For example, the two MS patients shown in Fig. 5 were symptom-free during the follow-up scans after the steroid treatments (Supplementary Materials); however. differential tractography still captures a substantial number of findings related to the primary visual pathways, and the decrease of the anisotropy diffusion correlates well with the severity of the initial clinical presentation. Similarly, by lowering the detection threshold, the ALS patient in Fig. 5 also shows substantial involvement in the posterior corpus callosum that connects to the occipital lobe. Although there is no clinical presentation of this patient associated with the findings, subclinical callosal damage for ALS patients is not uncommon (Filippini et al., 2010), and there are also studies suggested subclinical involvement of occipital lobe in the ALS patients (Loewe et al., 2017; Zhang et al., 2017). The ability to capture subclinical findings has a profound clinical implication. It suggests that differential tractography is sensitive enough to provide additional evaluation value on top of existing clinical scales and scores. This may facilitate the development of new treatment to target early subclinical change that may end up with irreversible damage.

### Limitations and possible pitfalls

There are limitations in differential tractography. Differential tractography only works on longitudinal scans of the same subject and only reflects the change of anisotropy within the time frame of the repeat scans. It cannot access track integrity in a cross-sectional setting, nor does it able to detect any abnormality or axonal injury prior to the baseline scans. Furthermore, differential tractography still has false results if the artifact also propagates coincidentally along a fiber pathway. The parallel imaging or eddy current artifact often gives rise to straight lines near the brain surface but may appear like a spuriously legitimate connection. Misalignment between baseline and follow-up scans can also generate a false result, and it can happen due to registration error or brain tissue shift after surgical intervention.

There are still other possible causes of false results. The subjects may have substantial movement in the follow-up scan, but not in the study or sham scan. There may be inconsistency in image acquisition between repeat scans such as changing the head coils or scanning protocol. Both scenarios can produce spurious findings, and thus a series of quality control is always needed to avoid these getting false results. Furthermore, the findings in differential tractography still need to be validated against neuroanatomy. Spurious findings often appear near the brain surface with odd trajectories (straight lines), while true findings tend to follow the trajectories of well-known neuroanatomical pathways. Prior neuroanatomy knowledge may help exclude false results from true findings.

There are also limitations in the reliability assessment. The sham setting in this study uses only one scan, and thus the calculated FDR only considered false results due to local random error (e.g. noise which randomly occurs at each imaging voxel) and does not include those due to systematic errors (errors that affects all imaging voxels simultaneously) such as subject movements, coil quality, or signal drift. To detect these systematic errors, we introduced four quality-control routines in the Material and Methods section to discard scans that had a substantial amount of systematic errors. Moreover, we did not acquire an additional sham scan in this study and used the “substitute sham” approach. The substitute sham approach can overestimate the number of false results due to a genuine increase of anisotropy (e.g. re-myelination after recovering from a neuronal injury). The consequence is that the FDR could be higher than the actual value, and the reliability of the findings could be underestimated, leading to a risk of missing meaningful findings.

Last, the diffusion MRI protocol in this study could be further optimized. Our result showed that high b-value played an important role in boosting the sensitivity of differential tractography, but the optimal value range still needs further investigation. Moreover, this study did not utilize the recent multi-band acquisition to reduce the scanning time, and future studies will utilize multi-band sequences to make differential tractography more feasible for clinical applications (e.g. 6-minutes grid-128 and 12-minute grid-256 acquisitions mentioned in the previous paragraphs).

### Clinical applications

Differential tractography can be used in differential diagnosis or prognostic evaluation after treatment or intervention. Neurologists can use it to differentiate the cause of a neurological disorder as patients with different neurological disorders will present distinctly different spatial patterns in their affected pathways. The location will provide a clue about the possible causes to resolve difficult clinical cases. This is otherwise not achievable in structural MRI unless a gross lesion or atrophy is visible in the late stage of neuronal injury. Another application of differential tractography is for evaluating an intervention or treatment. Differential tractography can provide an objective quantitation that is directly comparable across subjects and less susceptible to observer differences. It could minimize variance due to evaluator differences and increase effect size in comparison with the conventional evaluation conducted by patients or neurologists. This opens a gate for early treatments to restore subclinical injuries before those injuries accumulate to become a major functional deficit.

## Acknowledgments

The research reported in this publication was partly supported by NIMH of the National Institutes of Health under award number R56MH113634. The content is solely the responsibility of the authors and does not necessarily represent the official views of the National Institutes of Health. The MRI scans were partly supported by the Walter L. Copeland Fund of The Pittsburgh Foundation.

## Supplementary Materials

### Patient’s medical history during the scan interval

#### MS patients

Patient #1 was a 44-year-old female with acute onset of left ocular pain and pain with eye movements. She experienced a gradual decline in her visual acuity, and initial visual examination showed normal acuity in the right eye and decreased acuity in the left eye (20/70), which worsen to 20/400 in one week. She had a dense superior visual field defect in both superior quadrants that continued to worsen over a few days with gradual loss of the inferior visual fields. She was treated with IV steroids, and after the treatment, the ocular pain, pain with eye movement, and the visual field were improved. After 6 months of her initial presentation, her visual acuity returned to 20/20 both eyes and the visual fields were normal

Patient #2 was a 23-year-old female with the blurring of vision of the right eye, associated with ocular pain and pain with extraocular movements. Initial evaluation by ophthalmology noted visual acuity of 20/40 with enlargement of the blind spot on the Humphrey visual field. Within a week her symptoms progressed to a visual acuity of 20/125, decrease in the color perception. and a superior altitudinal visual field defect. She was treated with intravenous methylprednisolone according to the optic neuritis treatment trial. Her symptoms recovered within 10 days to visual acuity of 20/25 and normal color saturation.

#### HD patients

Patient #3 was a 60-year-old male. The age of onset was 56, and the CAG repeats were 42. During the scan interval, the UHDRS motor total was increased from 45 to 49, with most prominent symptoms of body bradykinesia on both sides.

Patient #4 was a 55-year-old female. The age of onset was 48, and the CAG repeats were 43. During the scan interval, the UHDRS motor total was increased from 53 to 64, with symptoms of dystonic contractures of the left upper extremity, dragging right foot with ambulation, and dystonia in the legs more obvious with gait (mostly on the right side), moderate bradykinesia with the finger taps (mostly on the left side).

#### ALS patient

Patient #5 was a 48-year-old male ALS patient (laboratory-supported probable ALS diagnosis) with upper extremity limb onset and predominant lower motor neuron involvement. His symptoms started about 30 months before the baseline scan with widespread fasciculations and left-hand weakness. There were subtle upper motor neuron signs in cervical segments. He developed lower extremity weakness 2 years after onset but never developed lower extremity upper motor neuron signs at the time of death that occurred 5 years after the onset of symptoms. He had no family history of neurological disease. The patient’s ALSFRS-R score decreased from 45 to 32 during the MRI scan interval (1 year apart).

#### Epilepsy patient

Patient #6 was a 51-year-old male with medically intractable epilepsy since the age of 17. His seizure frequency was weekly, and intracranial monitoring demonstrated that his seizure onset zone included the right mesial and neocortical temporal lobe. He underwent a standard right anterior temporal lobectomy. MRI scans were obtained before the surgery and one year after the surgery.

